# Hyper3D-lite: count-preserving representation auditing for long-read multi-contact genome data

**DOI:** 10.64898/2026.06.09.730992

**Authors:** Yubo Zhang

## Abstract

Long-read and single-molecule sequencing technologies are rapidly increasing molecule-level data, with platforms such as Oxford Nanopore, PacBio HiFi, and Roche sequencing-by-expansion advancing at different technology readiness levels. In the specific context of Pore-C and HiPore-C multi-contact chromatin-conformation assays, long-read multi-contact 3D genome assays preserve molecule-level contact context, but common downstream pairwise projections can expand one multi-contact molecule into many pair records. This creates a representation problem: apparent contact evidence can increase through the counting frame before biological interpretation begins. Hyper3D-lite addresses this problem as a representation-first audit tool for read-to-fragment-style long-read multi-contact inputs. It compares all-pair projection with CPB, a count-preserving statistical accounting reference point, and separates broad software outputs from conservative higher-order candidate calls.

## Introduction

Long-read multi-contact assays such as Pore-C and HiPore-C can retain concatemer-scale information that is not naturally represented by pairwise contact matrices alone [1,2]. This creates an analysis challenge: a single molecule with multiple contacted fragments can generate multiple pairwise records when expanded into all pairs. Pairwise projection is useful and familiar, but without explicit accounting it can obscure the relationship between molecule counts, projected pair counts, and higher-order software records.

Beyond these specific multi-contact assays, the broader long-read and single-molecule sequencing landscape motivates the need for representation-aware analysis. Oxford Nanopore sequencing has reached benchmark-scale throughput for structural variant detection, repeat expansion profiling, and transcript characterization in both research and clinical contexts [7]. PacBio HiFi long-read sequencing now supports cohort-scale structural variant discovery and resolves complex gene regions that are difficult to access with short reads [8,9]. Roche sequencing-by-expansion (SBX), as implemented in the development-stage AXELIOS platform, represents an emerging single-molecule sequencing approach with distinct chemistry [10,11]. These platforms differ in chemistry, error profile, and read length, but they share a common feature: they produce multi-fragment molecule-level data when applied to chromatin-conformation assays. It is important to distinguish ordinary long-read sequencing applications (genome assembly, variant detection, transcriptomics) from the specific use of long reads in Pore-C and HiPore-C multi-contact chromatin-conformation assays, where concatemer preservation creates the representation challenge that needs to be addressed. As platform throughput increases, the risk of conflating projected pairwise counts with independent biological observations grows, reinforcing the need for explicit representation auditing before mechanism interpretation.

Hyper3D-lite is designed as a representation-first audit layer for this setting. Rather than beginning with biological interpretation, it quantifies how much all-pair projection expands the input relative to CPB, a count-preserving accounting baseline. CPB is a statistical accounting reference point, not a simulation of polymer physics, ligation kinetics, chromatin dynamics, or nuclear mechanics. This distinction matters because recent 3D genomics reviews emphasize that assay design, resolution, and representation choices shape downstream interpretation [3]. We therefore frame the present study. The HBB/HBB-LCR region provides a biologically interpretable testbed for this method-to-mechanism bridge. Prior beta-globin perturbation and looping studies demonstrate that LCR-promoter contact configurations can be experimentally manipulated in appropriate systems [4–6]. These studies motivate HBB/HBB-LCR as a meaningful regulatory locus.

## Results

### Hyper3D-lite quantifies pairwise projection inflation using CPB

Long-read multi-contact data can preserve molecule-level context, but downstream all-pair expansion can produce many pair records from one read or concatemer [1,2]. Hyper3D-lite treats this as a representation problem. For each read-to-fragment-style input, the method reports projected pair records, higher-order software records when available, and an all-pair vs CPB inflation factor (Fig. 1). CPB is used as a count-preserving statistical accounting baseline. It is not a physical null model, not a polymer simulation, and not active noise filtering. This accounting layer makes the counting frame explicit before any biological claims are considered. In the current full-paper scope, CPB supports method-level interpretation only.

**Figure 1.**
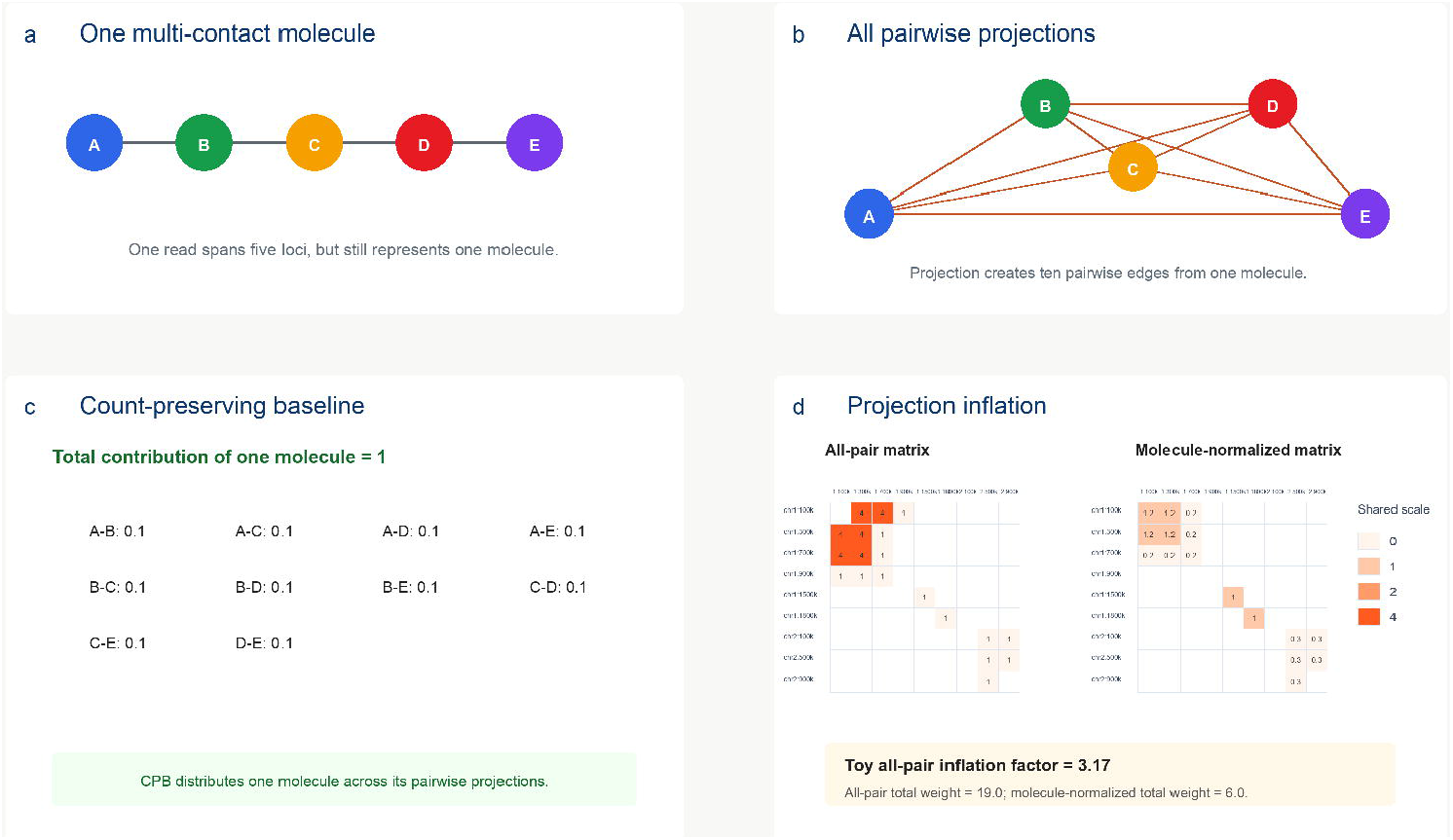
Count-preserving representation audit of pairwise projection inflation. **a**, A multi-segment long-read contact can encode more than two genomic fragments. **b**, Naive all-pair projection expands this representation into multiple pairwise contacts. **c**, The Count-Preserving Baseline (CPB) provides a statistical accounting reference point for preserving read-level support. **d**, Hyper3D-lite reports projection inflation relative to CPB. CPB is not a polymer-physics or chromatin-dynamics model.

### Toy examples demonstrate conservative higher-order candidate handling

Toy examples establish expected software behavior under controlled inputs and provide a minimal check that higher-order candidate outputs remain conservative. Published high-order Pore-C analyses show that higher-order candidate detection requires explicit statistical framing and background assumptions [12]. In Hyper3D-lite v1 drafting, toy examples are used for representation behavior and conservative candidate handling; they are not biological discovery experiments (Fig. 2). This conservative stance carries through the real-data sections. Candidate cliques are reported as exploratory software outputs unless they pass explicit significance and validation gates. Significant cliques remain the relevant conservative output class for any later biological interpretation, and no validated regulatory clique claim is made here.

**Figure 2.**
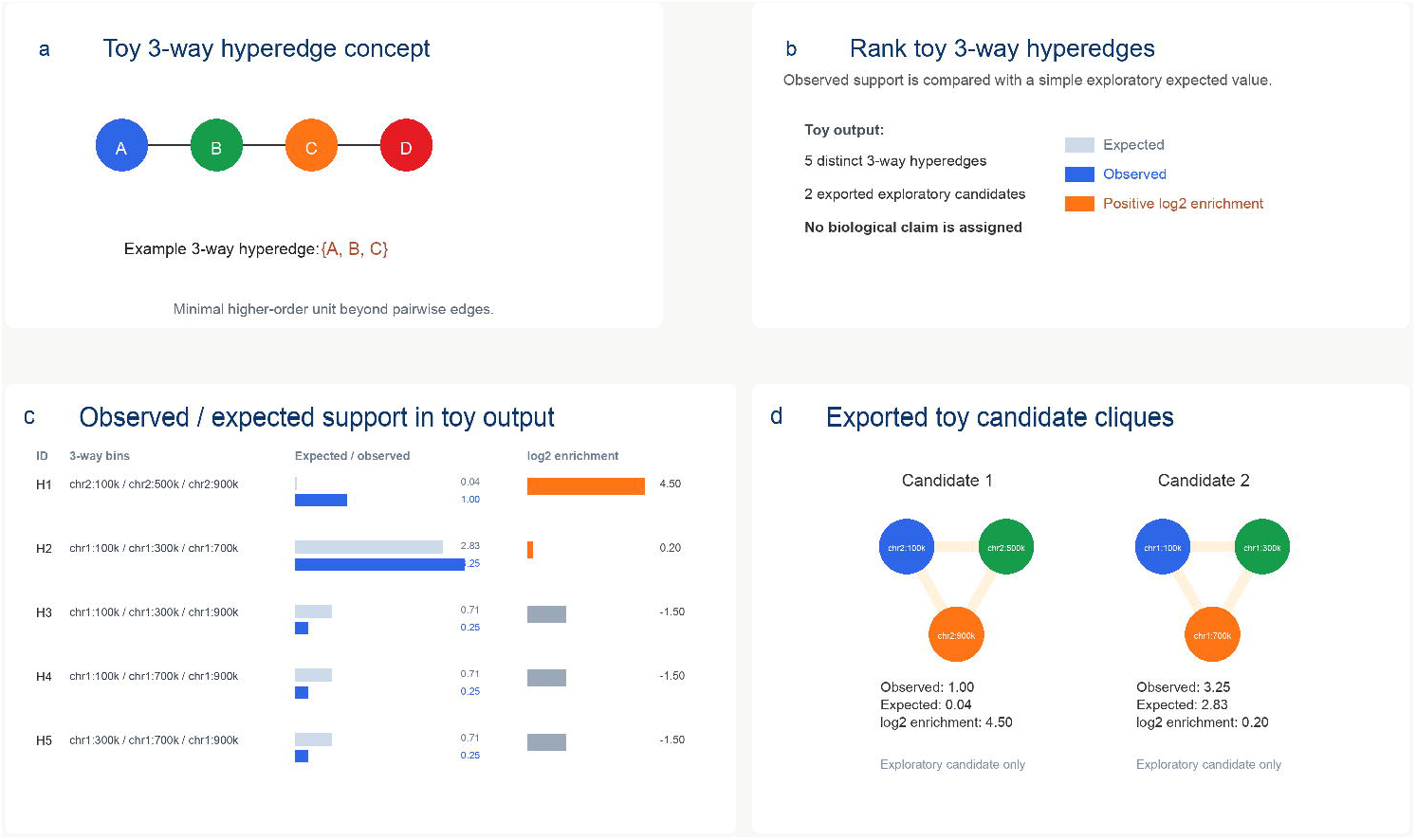
Toy-level higher-order candidate handling. **a**, Toy multi-segment reads illustrate how Hyper3D-lite represents higher-order contacts. **b**, Three-way hyperedges are generated as software-level representations. **c**, Candidate filtering separates recurrent support from dispersed singleton support. **d**, Output classes are used for representation auditing. Toy examples are illustrative only and are not interpreted as biological interactions.

### K562_FC3 HBB/HBB-LCR locus-focused analysis

K562_FC3 HBB/HBB-LCR locus-focused full-read-context input was derived from public GSE202539/HiPore-C data [1,13]. The locus window was chr11:5225463-5304186 in hg38 / GRCh38, using a PAF-derived, BED-like 0-based half-open coordinate convention. The extraction retained full-read context for locus-touching reads (Fig. 3).

**Figure 3.**
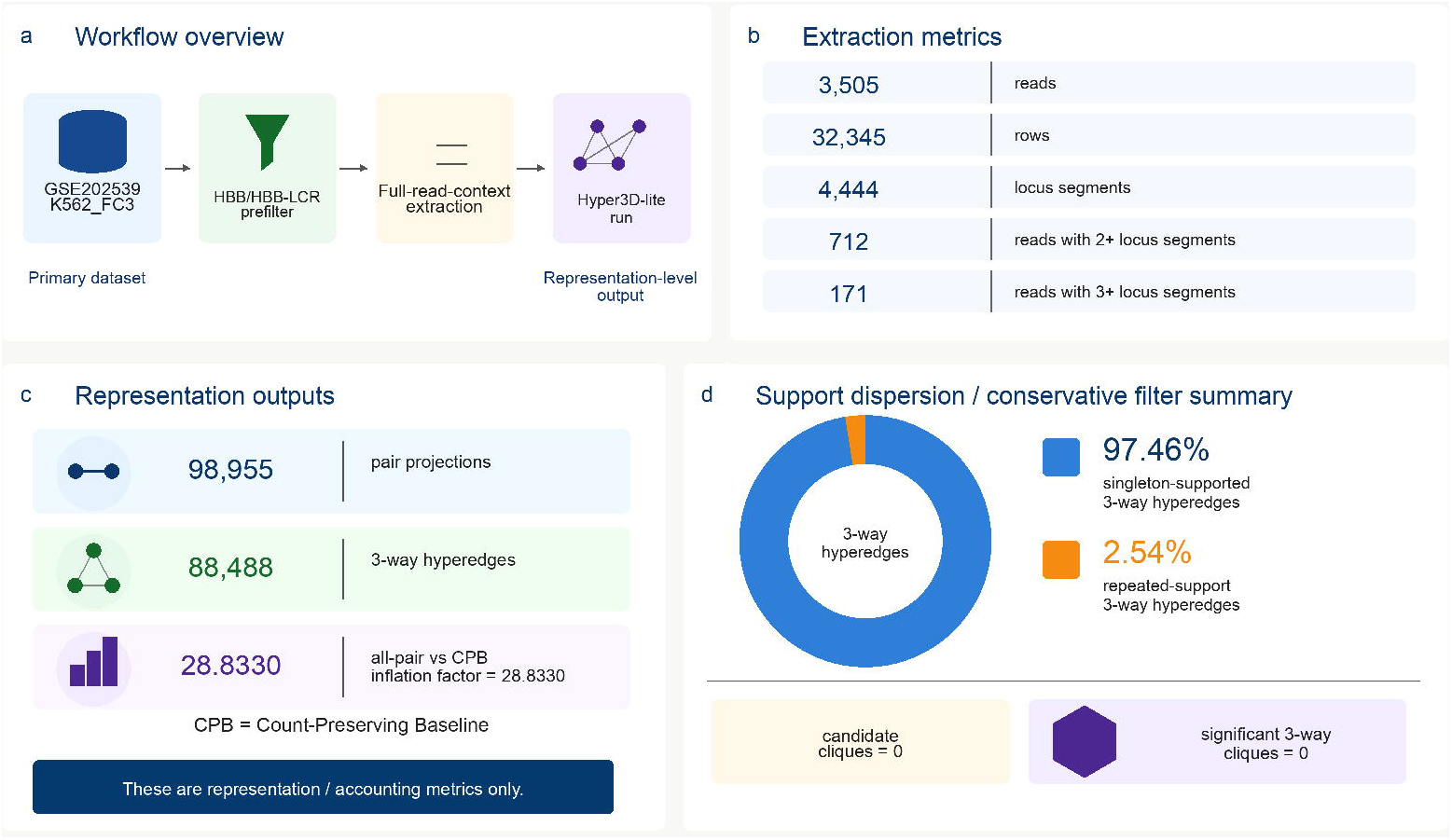
K562_FC3 HBB/HBB-LCR locus-focused representation audit. **a**, K562_FC3 reads from GSE202539 were processed through HBB/HBB-LCR prefiltering, full-read-context extraction and Hyper3D-lite analysis. **b**, The locus-focused input contained 3,505 extracted reads, 32,345 rows and 4,444 locus segments. **c**, Hyper3D-lite produced 98,955 pair projections and 88,488 three-way hyperedges, with an all-pair versus CPB inflation factor of 28.83. **d**, Support was dominated by singleton-supported three-way hyperedges (97.46%), with no significant three-way cliques detected. This is a method-focused representation audit, not evidence for an HBB/HBB-LCR mechanism.

The K562_FC3 run included 3,505 extracted reads and 32,345 extracted rows, including 4,444 locus segments, 712 reads with 2+ locus segments, and 171 reads with 3+ locus segments. Hyper3D-lite generated 98,955 pair projection rows and 88,488 3-way hyperedge software records. The all-pair vs CPB inflation factor was 28.83.

Candidate filtering was conservative. K562_FC3 produced candidate cliques = 0 and significant 3-way cliques = 0. The 3-way support distribution was dominated by singleton-supported hyperedges, with 97.46% singleton-supported and 2.54% repeated-support hyperedges. This supports a representation-level method result and a zero-candidate diagnostic explanation based on support dispersion and conservative filtering. It does not support an HBB biological mechanism claim, K562 erythroid mechanism claim, enhancer-promoter validation, or validated regulatory clique claim [14] (Fig. 4).

**Figure 4.**
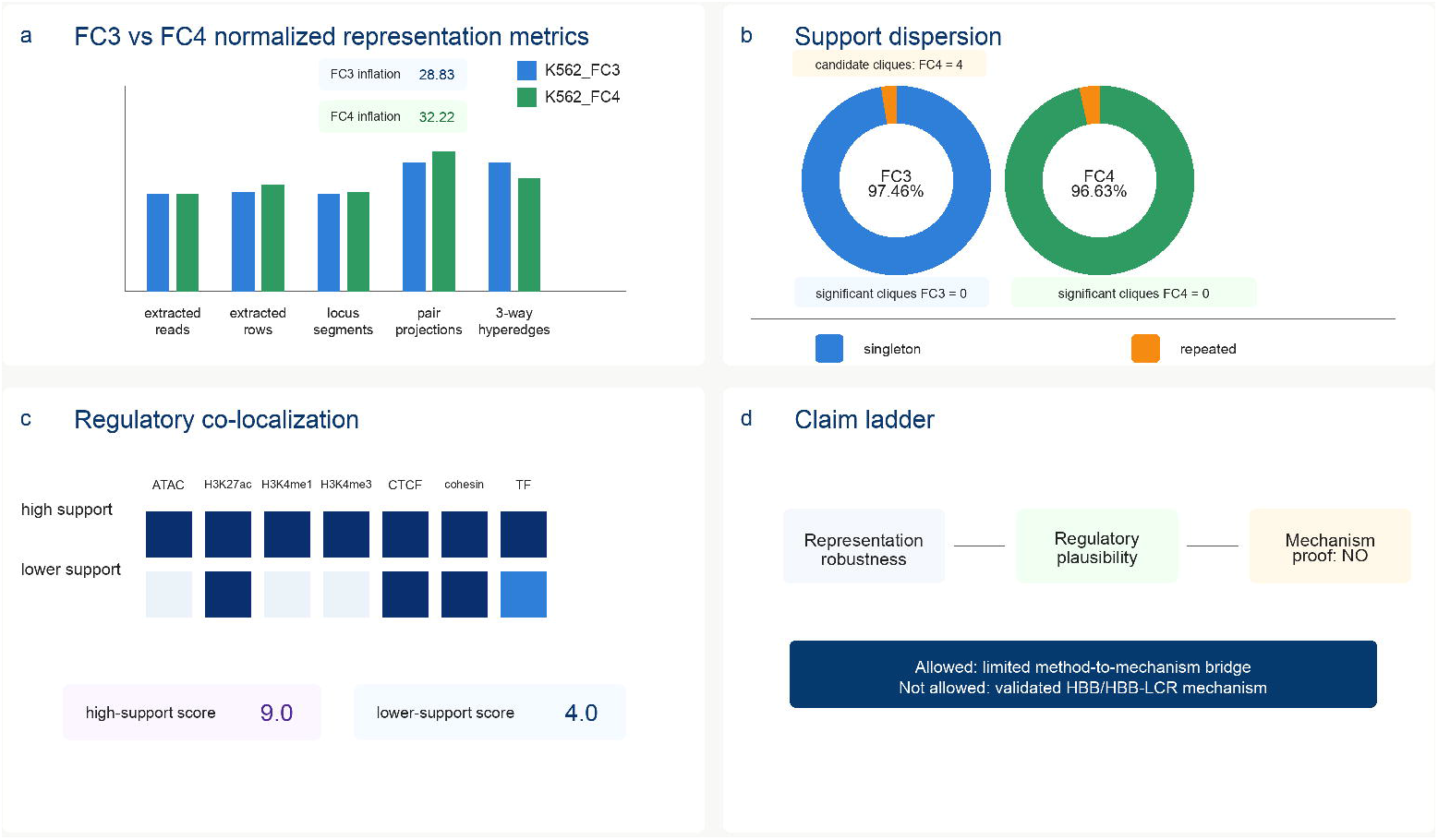
K562_FC3/K562_FC4 representation robustness and regulatory co-localization support. **a**, Depth-aware comparison of K562_FC3 and K562_FC4 representation-level metrics at the HBB/HBB-LCR locus. **b**, Both flowcells showed high singleton-supported three-way hyperedge fractions and no significant three-way cliques; FC4 produced four exploratory candidate cliques, none of which were significant. **c**, ENCODE-derived K562 regulatory annotations were summarized as co-localizing annotation classes, not as functional or causal scores. **d**, The evidence supports representation robustness and regulatory plausibility, but not enhancer-promoter causality, a validated regulatory clique or a proven HBB/HBB-LCR mechanism. Evidence-summary figure only; not a biological mechanism claim.

### Concordant representation-level robustness analysis

K562_FC4 was acquired as one additional K562 flowcell from the same public GSE202539/HiPore-C source context [1,13]. FC4 completed schema sanity, HBB/HBB-LCR full-read-context extraction, Hyper3D-lite run, overlap/redundancy diagnostic, and FC3-vs-FC4 comparison.

K562_FC4 streamed 87,997,619 rows, extracted 1,810 reads and 18,744 rows, identified 2,356 locus segments, and included 110 reads with 3+ locus segments.

Hyper3D-lite produced 56,771 pair projection rows and 40,582 3-way hyperedges. The all-pair vs CPB inflation factor was 32.22.

FC3 and FC4 show concordant representation-level behavior. FC3 and FC4 both show strong all-pair vs CPB inflation, with inflation factors of 28.83 and 32.22. Both flowcells also show singleton-dominated 3-way hyperedge support dispersion: FC3 singleton-supported fraction = 97.46%, and FC4 singleton-supported fraction = 96.99%. Significant cliques remain zero in both flowcells. FC4 produced 4 candidate cliques, but these are exploratory software-level candidates only and are not interpreted as biological interactions.

The normalized comparison supports representation-level robustness across K562 flowcells. It does not establish independent biological replicate support, enhancer-promoter causality, or an HBB/HBB-LCR regulatory mechanism. High-order contact studies provide useful representation frameworks, but candidate calls require explicit statistical and biological validation before mechanism interpretation [2,12].

### Annotation-only and multi-omics overlap support regulatory plausibility

Annotation-only analysis used a finalized manifest that avoided unverified HS/promoter coordinate assumptions. Exact HS element coordinates were not invented. The executable annotation layer was therefore limited to verified HBB/HBB-LCR union-level coordinates and project-derived representation bins.

Nine ENCODE K562 GRCh38 processed BED peak files were acquired and used for multi-omics overlap. ENCODE K562 accessions provide public regulatory annotation provenance for chromatin activity, cohesin/architecture, boundary-factor, and transcription factor context [15–18]. The support-tier summary found co-localization between evaluated HBB/HBB-LCR representation bins and regulatory annotations. In the current summary, the high-support tier had mean regulatory feature score 9.0 across one evaluated item, while the lower-support tier had mean regulatory feature score 4.0 across one evaluated item. These overlaps support regulatory plausibility for using HBB/HBB-LCR as a method-to-mechanism bridge [19–21].

## Discussion

Hyper3D-lite starts from a representation problem. Long-read multi-contact molecules can carry more context than pairwise records, but the act of pairwise expansion changes the counting frame. By reporting all-pair vs CPB inflation and separating exploratory software outputs from conservative candidate calls, Hyper3D-lite provides a method-level audit layer before mechanism interpretation [1,3].

HBB/HBB-LCR is useful here because it is biologically interpretable and historically well studied. Possible mechanisms include LCR-promoter looping, multi-way enhancer/ promoter hub organization, CTCF/cohesin-associated architecture, and active chromatin or erythroid transcription-factor-associated contact enrichment. Prior forced-looping, CLOuD9, LDB1/TAL1, and multi-way globin literature make these reasonable hypothesis classes [4–6,22]. The FC3/FC4 comparison strengthens representation-level robustness. Both K562 flowcells show strong all-pair vs CPB inflation and singleton-dominated 3-way support dispersion, and both have zero significant cliques. This reduces concern that the representation pattern is a single-flowcell artifact. It does not convert flowcell consistency into independent biological replicate validation or mechanism proof.

The broader long-read sequencing landscape reinforces the methodological importance of representation auditing. Nanopore, PacBio HiFi, and Roche SBX/AXELIOS represent distinct approaches to single-molecule sequencing with different chemistry, error profiles, throughput characteristics, and technology readiness levels [7–10]. Despite these differences, they share a common downstream challenge when used for multi-contact chromatin-conformation assays: preserving molecule-level context through the analytical pipeline. Pore-C and HiPore-C are the specific assays where this challenge arises most directly, because each multi-fragment concatemer can generate many pairwise projections. As platform capabilities scale, so does the volume of multi-contact data, making projection-aware representation auditing increasingly important. Hyper3D-lite provides this audit layer before biological interpretation begins, separating representation inflation from biological signal. Platform progress motivates the importance of the method; it does not validate the specific HBB/HBB-LCR observation or any biological mechanism claim. ENCODE co-localization adds regulatory plausibility. K562 regulatory peaks overlap evaluated HBB/HBB-LCR representation bins, consistent with the region’s use as a biologically interpretable testbed. But co-localization is correlative. It does not prove enhancer-promoter contact function, establish causality, or validate FC4 candidate cliques as biological interactions [14].

Third-party evidence therefore plays a bounded role. It supports the assay rationale, the HBB/HBB-LCR testbed rationale, ENCODE annotation provenance, and the methodological need for background models in enrichment analysis. It does not validate the project-specific mechanism. To upgrade the mechanism claim, it would need independent biological replicates, orthogonal 3D genome assays, perturbation evidence, RNA or expression readouts, preregistered background enrichment, and further works.

## Methods

### Hyper3D-lite representation workflow

Hyper3D-lite takes read-to-fragment-style long-read multi-contact inputs and reports representation-level outputs, including pair projections, higher-order hyperedge software records where supported, candidate clique outputs, and QC summaries. The workflow is intended to audit representation behavior before biological interpretation.

### CPB accounting baseline

CPB is a count-preserving statistical accounting baseline. It provides a reference point for comparing all-pair projection against a molecule-preserving counting frame. CPB is not a physical null model, not a polymer simulation, and not an active denoising method.

### K562_FC3 and K562_FC4 provenance

K562_FC3 and K562_FC4 inputs were derived from public GSE202539/HiPore-C source data [1,13]. GSE202539 is cited as a GEO accession-based source; no article DOI is invented for the accession. Raw GSE202539 data are not redistributed in the project package.

### HBB/HBB-LCR locus definition

The HBB/HBB-LCR planning union was defined as chr11:5225463-5304186 in hg38 / GRCh38. Coordinates follow a PAF-derived, BED-like 0-based half-open convention.

### Full-read-context extraction

The extraction procedure identified reads touching the HBB/HBB-LCR planning union and retained full-read context for those reads. This preserves non-locus segments present on locus-touching reads, allowing representation-level context to be audited rather than trimming each read to the locus interval only.

### Hyper3D-lite run and diagnostics

K562_FC3 and K562_FC4 locus-focused full-read-context inputs were processed with the current Hyper3D-lite run workflow. Run reports collected input rows, unique reads/ concatemers, pair projection rows, 3-way hyperedges, all-pair vs CPB inflation factor, candidate clique counts, significant clique counts, runtime, and output files. Overlap/ redundancy diagnostics summarized supporting-read distributions and candidate-output behavior.

### FC3/FC4 normalized metric comparison

FC3/FC4 consistency was assessed using raw and normalized metrics. Extraction metrics were normalized per 1M streamed rows, and downstream representation outputs were normalized per 1k extracted reads where appropriate. These normalizations reduce depth-driven confusion but do not convert flowcells into biological replicates.

### Annotation-only overlap

Annotation-only analysis used a finalized HBB/HBB-LCR manifest. Coordinates were included only when sufficiently verified for the project boundary. Exact HS element coordinates were not invented. The executable overlap analysis was limited to safe HBB/HBB-LCR union-level and representation-bin features.

### ENCODE K562 multi-omics overlap

Nine ENCODE K562 GRCh38 processed BED peak files were acquired for multi-omics overlap. Raw FASTQ/BAM files were not downloaded. Overlap outputs included a regulatory feature support matrix and support-tier summary. These outputs were interpreted as co-localization plausibility only.

## Supporting information

K562_HBB_LCR_method_metrics

overlap_redundancy_result

hyperedge_support_distribution

## Data and Code Availability

This draft uses the current project development package and local outputs. Public release decisions remain separate from this draft. GSE202539 raw data are public source data and are cited from the original GEO/HiPore-C source context; they are not redistributed here. GitHub and DOI assignment are not created in this drafting step.

Local development paths are not public accessions. Derived small method-level outputs, figures, and tables can be prepared for a later full-paper submission package only after user approval.

## Supplementary Information

Supplementary materials include K562_HBB_LCR_method_metrics, overlap_redundancy_result, and hyperedge_support_distribution.

